# Neutralization of recombinant RBD-subunit vaccine ZF2001-elicited antisera to SARS-CoV-2 variants including Delta

**DOI:** 10.1101/2021.07.15.452504

**Authors:** Xin Zhao, Anqi Zheng, Dedong Li, Rong Zhang, Huan Sun, Qihui Wang, George F. Gao, Pengcheng Han, Lianpan Dai

## Abstract

SARS-CoV-2 variants brought new waves of infection worldwide. In particular, Delta variant (B.1.617.2 lineage) has become predominant in many countries. These variants raised the concern for their potential immune escape to the currently approved vaccines. ZF2001 is a subunit vaccine received emergency use authorization (EUA) in both China and Uzbekistan, with more than 100-million doses administrated with a three-dose regimen. The tandem-repeat dimer of SARS-CoV-2 spike protein receptor binding domain (RBD) was used as the antigen. In this work, we evaluated the neutralization of ZF2001-elicited antisera to SARS-CoV-2 variants including all four variants of concern (Alpha, Beta, Gamma and Delta) and other three variants of interest (Epsilon, Eta and Kappa) by pseudovirus-based assay. We found antisera preserved majority of the neutralizing activity against these variants. E484K/Q substitution is the key mutation to reduce the RBD-elicited sera neutralization. Moreover, ZF2001-elicited sera with a prolonged intervals between the second and third dose enhanced the neutralizing titers and resilience to SARS-CoV-2 variants.

---

SARS-CoV-2 variants are continually emerging and becoming the circulating strains in many countries. Several highly transmissible variants of concern (VOCs) showed altered pathogenicity and become dominant worldwide, including B.1.1.7 (Alpha), B.1.351 (Beta), P.1 (Gamma) and B.1.617.2 (Delta) lineages (www.who.int). Some variants were reported to escape the immunity acquired by natural infection or vaccination, bringing a global concern for the effectiveness of the currently approved vaccines^1^. In particular, the recently emerged Delta variant have caused another wave of infection worldwide^2–5^. The effect of the mRNA vaccines (mRNA-1273 and BNT162b2) and the adenovirus vector-based vaccine ChAdOx1 against Delta variants have been evaluated in both serum neutralization and real-world protection^4–11^. These vaccines showed various reduction of neutralization against Delta variant ranging from 1.4- to 9-fold based on different methods^4–9^, and substantial reduction in real-world protection^10,11^.

ZF2001 is a protein subunit vaccine, currently in phase 3 clinical trials, using tandem-repeat SARS-CoV-2 spike receptor-binding domain (RBD) dimer as the antigen^12,13^. ZF2001 has received emergency use authorization (EUA) in both China and Uzbekistan since March, 2021 and is rolling out for vaccination with a three-dose regimen. ZF2001-elicited serum showed a 1.6-2.8-fold reduction in neutralization of B.1.351 variant^14,15^. Here, we used the vesicular stomatitis virus (VSV)-based pseudotyped virus expressing SARS-CoV-2 spike to test the neutralization activity of ZF2001 to a panel of variants, including all four VOCs: Alpha (B.1.1.7 lineage), Beta (B.1.351 lineage), Gamma (P.1 lineage) and Delta (B.1.617.2 lineage). Another three important variants of interest (VOIs), Epsilon (B.1.429 lineage), Eta (B.1.525 lineage) and Kappa (B.1.617.1 lineages), were also included. SARS-CoV-2 Wild type strain (Wuhan-1 reference strain) was tested for comparison^16,17^. The mutation site of each variant in spike was listed in Supplementary Fig. 1.

Sera from 28 candidates (13 males and 15 females) receiving full vaccination of 3-dose ZF2001 were collected to measure the 50% pseudovirus neutralization titer (pVNT_50_). Detailed information regarding the vaccination and blood sampling time intervals were summarized in Table S1. We found all 28 serum samples efficiently neutralized pseudotyped virus expressing wild-type spike (Wuhan-1 reference strain), with the pVNT_50_ higher than 1:20. We found the neutralizing titer did not decline, but slightly increased (1.1 fold to WT, P>0.05), against either D614G- or B.1.1.7 (N501Y mutation in RBD)-spike pseudovirus (Fig.1A, B and Fig.S1-2). However, variant with single mutation at L452R in RBD (B.1.429 [Epsilon]-spike) or double mutations at both L452R and T478K in RBD (B.1.617.2 [Delta]-spike) showed roughly equivalent sensitivity to ZF2001-elicited antisera as compared with pseudovirus expressing WT-spike (−1.1 and −1.2 fold to WT, respectively; p>0.05) (Fig.1A, E, H and Fig.S1-2). Therefore, ZF2001 preserved the neutralizing activity against the newly emerging but highly transmissible Delta variant. In contrast, the variants with E484K/Q substitution in RBD showed more pronounced reduction in sensitivity (B.1.351 [Beta]-spike, −1.8 fold, p=0.0071; P.1 [Gamma]-spike, −1.5 fold, p=0.0505; B.1.525 [Eta]-spike, −2.0 fold, p=0.0021; B.1.617.1 [Kappa] spike, −2.1 fold, p<0.0001) (Fig.1A, C, D, F, G and Fig.S1-2), which is consistent with what has been reported for ZF2001-elicited antisera against authentic B.1.351 lineage virus^14,15^. Unlike other approved vaccines targeting whole virus or full spike protein, ZF2001 only targets RBD^12,18^. Our result in sera level is consistent with the previous finding in monoclonal antibody level that RBD-targeting antibodies are resilient to SARS-CoV-2 variants because of a more dispersed neutralizing epitopes in RBD compared with NTD^19^.

**Figure 1:**
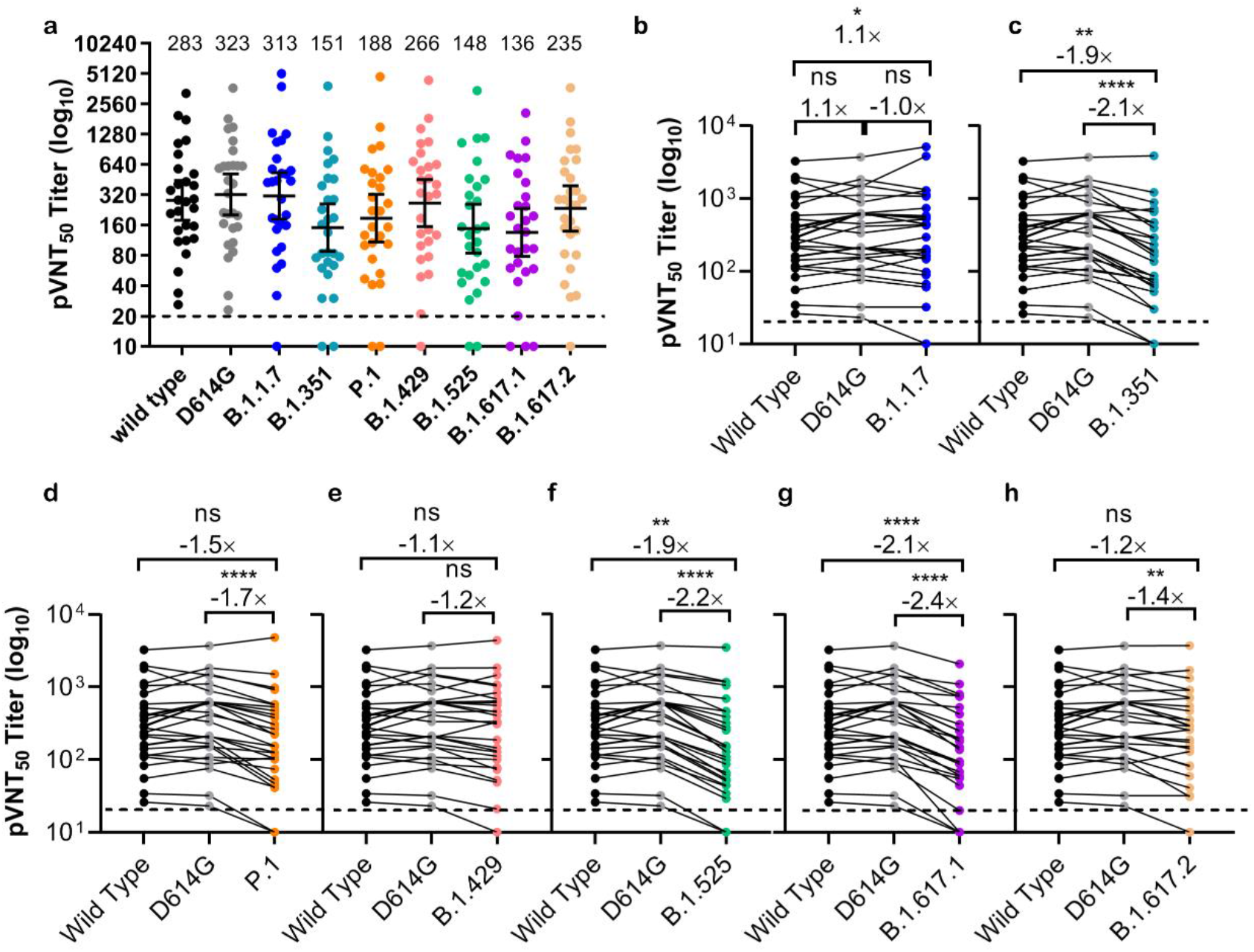
Pseudotyped virus neutralization of SARS-CoV-2 variants in ZF2001-elicited serum samples. **a:** 50% pseudovirus neutralization titer (pVNT_50_) against VSV-expressing wild type spike or eight spike variants of SARS-CoV-2 in sera from 28 volunteers receiving three doses of ZF2001 vaccination. The variants including the parental variant D614G, four VOCs (B.1.1.7, B.1.351, P.1 and B.1.617.2 lineages), and three variants of interest (B.1.429, B.1.525 and B.1.617.1 lineages) listed by WHO. The geometric mean titer (GMT) was marked on top of each column and lined with 95% confidential interval (CI) shown as horizontal bars. The dashed line indicates the lower limit of detection. The GMT lower than 20 were considered negative and calculated as 10 in the statistical analysis. **b-h** shows the pVNT_50_ titer against each of the seven variants compared with wild type and D614G. Fold change and significance compared with both wild type and D614G were shown in each panel. (*, p<0.05; **, p<0.01; **, p<0.001; ****, p<0.0001). Neutralization values between different variants were analyzed with two-tailed Wilcoxon matched-pairs signed rank test. pVNT_50_ of each sample is tested by two repeats.

Further, the volunteers with an extended interval between the second and third doses (0-1-(4-6) regimen) showed higher neutralizing activity and resilience to variants than those with shorter interval (0-1-2 regimen) (Fig.2), which is consistent with a prior work for neutralization of B.1.351 lineage virus by the ZF2001-elicited antisera^15^. In the 0-1-2 group, the antisera neutralization titers were significant reduced against six of the seven variants except B.1.1.7 (Figs.2A and 2C). In contrast, in the 0-1-(4-6) group, significant reduction of neutralization titer was only detected against the B.1.617.1-spike lineage virus (double mutations at L452R and E484Q in RBD) (Fig.2A, C and Fig. S1). The better performance of the 0-1-(4-6) regimen is likely due to the benefit from the prolonged antibody maturation in individuals^14^. Our data are consistent with the fact that the 0-1-6 regimen has been widely used for other subunit vaccines, such as Hepatitis B virus vaccine, and provide guidance to further optimize the vaccination regimen.

**Figure 2:**
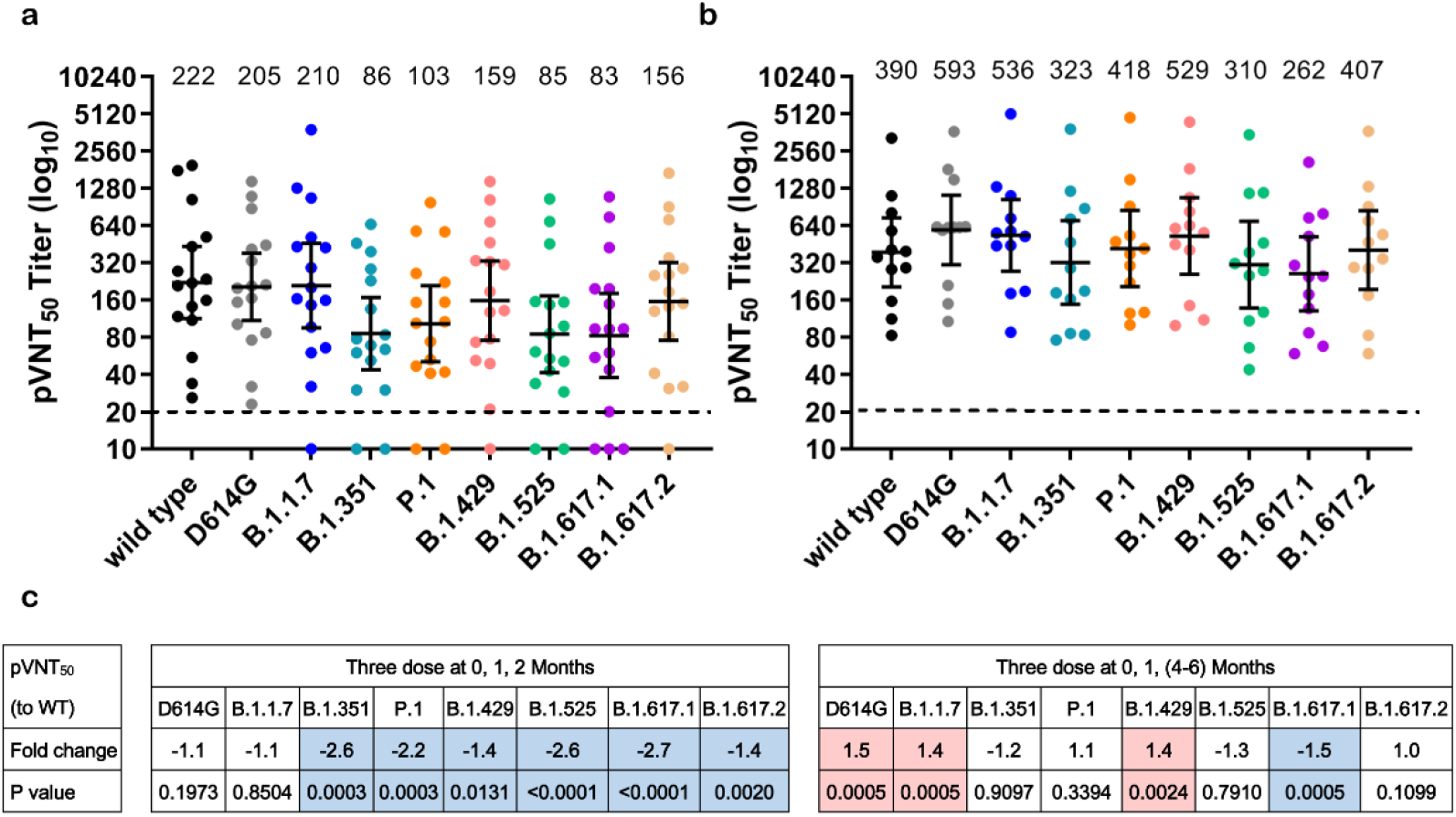
Pseudotyped virus neutralization of SARS-CoV-2 variants in ZF2001-elicited serum samples from people received three dosed in different vaccination regimen. **a:** The pVNT_50_ against variants in volunteers (n=16) with three dose of ZF2001 vaccination at 0, 1, 2 Months. **b**: The pVNT_50_ against variants in volunteers (n=12) with three doses at 0, 1, 4-6 Months. The geometric mean titer (GMT) was marked on top of each column and lined with 95% confidential interval (CI) shown as horizontal bars. The dashed line indicates the lower limit of detection. The GMT lower than 20 were considered negative and calculated as 10 in the statistical analysis. **c:** Summarization the fold-change and p value of the pVNT_50_ for the variants from the wild type. Background color in pink indicates significant GMT increase and in light blue indicates significant GMT decrease, compare with the GMT to the wild type. White means no significant change. Neutralization values between different variants were analyzed with two-tailed Wilcoxon matched-pairs signed rank test. pVNT_50_ of each sample is tested by two repeats.

In this work, we provide preliminary evidence of the approved RBD-based protein subunit vaccine for its neutralization profile to SARS-CoV-2 variants. Given the positive correlation between neutralizing titer and protection efficacy^20^, the sera neutralization variation between variants should be taken into account for the vaccine effectiveness. The high susceptibility of these newly emerged variants to ZF2001 vaccine supports the current mass immunization to build a herd immunity. However, the vaccine effectiveness against these variants must be validated by phase 3 clinical trials and real-world evidences.

## Acknowledgments

We thank all the volunteers for providing blood samples. This work was supported by the intramural special grant for SARS-CoV-2 research from the Chinese Academy of Sciences; the Strategic Priority Research Program of the Chinese Academy of Sciences (XDB29010202) to G.F.G. X.Z. is supported by Beijing Nova program of Science and Technology (Z191100001119030), and Youth Innovation Promotion Association of the CAS (20200920). L.D. is supported by Youth Innovation Promotion Association of the CAS (2018113).

## Author contributions

GFG, LD and PH conceived and designed the study. XZ, LD, PH and QW designed, and coordinated the experiments. XZ, AZ, DL and RZ performed experiments. HS recruited volunteers and coordinated the blood samples. XZ and AZ analyzed the data. GFG, XZ and LD drafted and revised the manuscript. All authors reviewed and approved the final manuscript.

