## Supplemental Material for "Neutralization of recombinant RBD-subunit vaccine ZF2001-elicited antisera to SARS-CoV-2 variants including Delta"

**Contents** **Page**

### Materials and Methods

#### Serum samples

The 28 serum samples were collected from vaccine recipients after 3 dose of 25 μg ZF2001 vaccine on 0, 1, 2 Months (16 vaccinees) or 0, 1, (4-6) Months (12 vaccinees)[1,2]. The participants including 13 males and 15 Females. Detailed information were provide in Table S1. All candidates signed the written informed consent.

#### Pseudotyped virus neutralization assay

The construction of VSV-ΔG-GFP based SARS-CoV-2 pseudotyped virus was mentioned in previous work with slight modifications[3-6]. The codon optimized SARS-CoV-2 wild type (Wuhan-1 reference strain) and variants spike protein (with mutations shown in Fig.S1) with an 18 amino acid truncation at the C-terminal were constructed into the pCAGGS vector. 30 μg of the construct was transfected into HEK 293T cells. VSV-ΔG-G-GFP pseudovirus were added 24 h after the transfection and removed after 1 h incubation. Medium were changed into fresh complete DMEM medium with anti-VSV-G antibody (I1HybridomaATCC® CRL2700™). Supernatants were collected after another 30 h incubation, passed through a 0.45 μm filter (Millipore, Cat#SLHP033RB), aliquoted and stored at -80 °C.

For the neutralization assay, the heat-inactivated (56 °C for 30min) serum sample from vaccinees were 2-fold serial diluted started from 1:20. 50 μL of the serially diluted sera were incubated with 50 μL of each pseudoviruses at 1000 transducing units (TU) at 37 °C for 1 h, and added onto pre-plated Vero cells (ATCC CCL81) in 96 well plate. The TU numbers were calculated after a 15 h incubation on a CQ1 confocal image cytometer (Yokogawa) [3].

**Data analyzing and statistical analysis**

The TU numbers were calculated by CQ1 software and statistical analyses were performed by GraphPad Prism 8.0 (GraphPad Software Inc.) for all experiments. $pVNT_{50}$ were determined by a non-linear regression (Fig. S2). $pVNT_{50}$ below the lower limit of detection (<20) were recorded as 10 in the geometric mean calculation. Neutralization values between different strains were compared using a two-tailed Wilcoxon matched-pairs signed rank test.

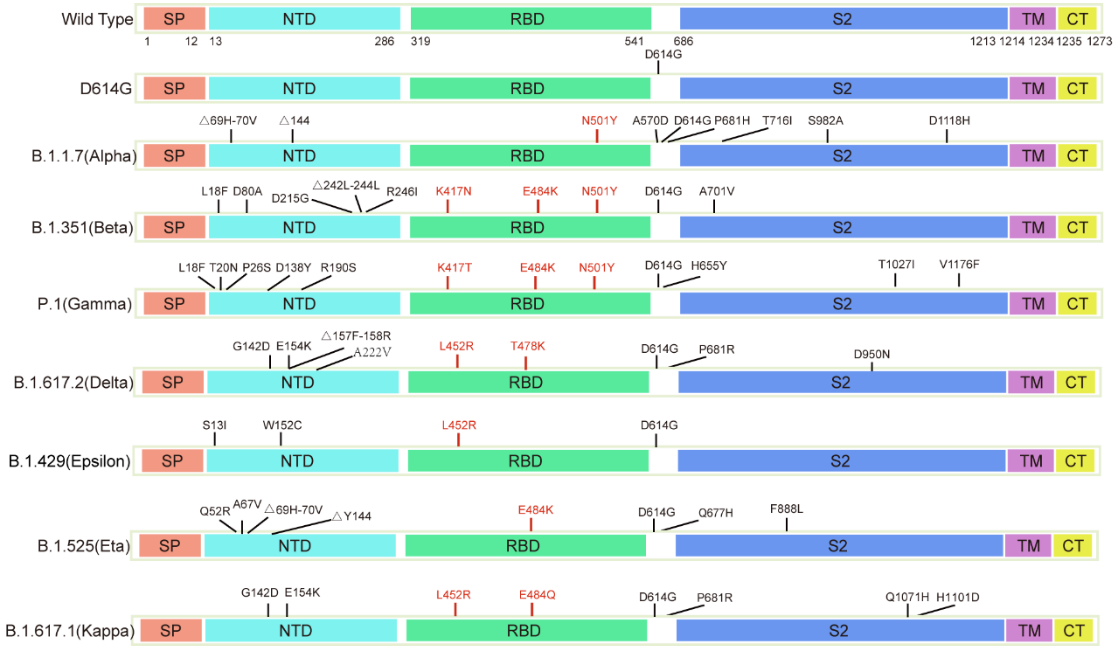

**Figure S1: Schematic representation of SARS-CoV-2 Spike proteins including wild type (WT) and eight variants.** The mutation sites of the variants were indicated, with the mutations on RBD highlighted in red. (SP: signal peptide, NTD: N-terminal domain, RBD: receptor-binding domain, TM: transmembrane domain, CT: C-terminal cytoplasmic domain).

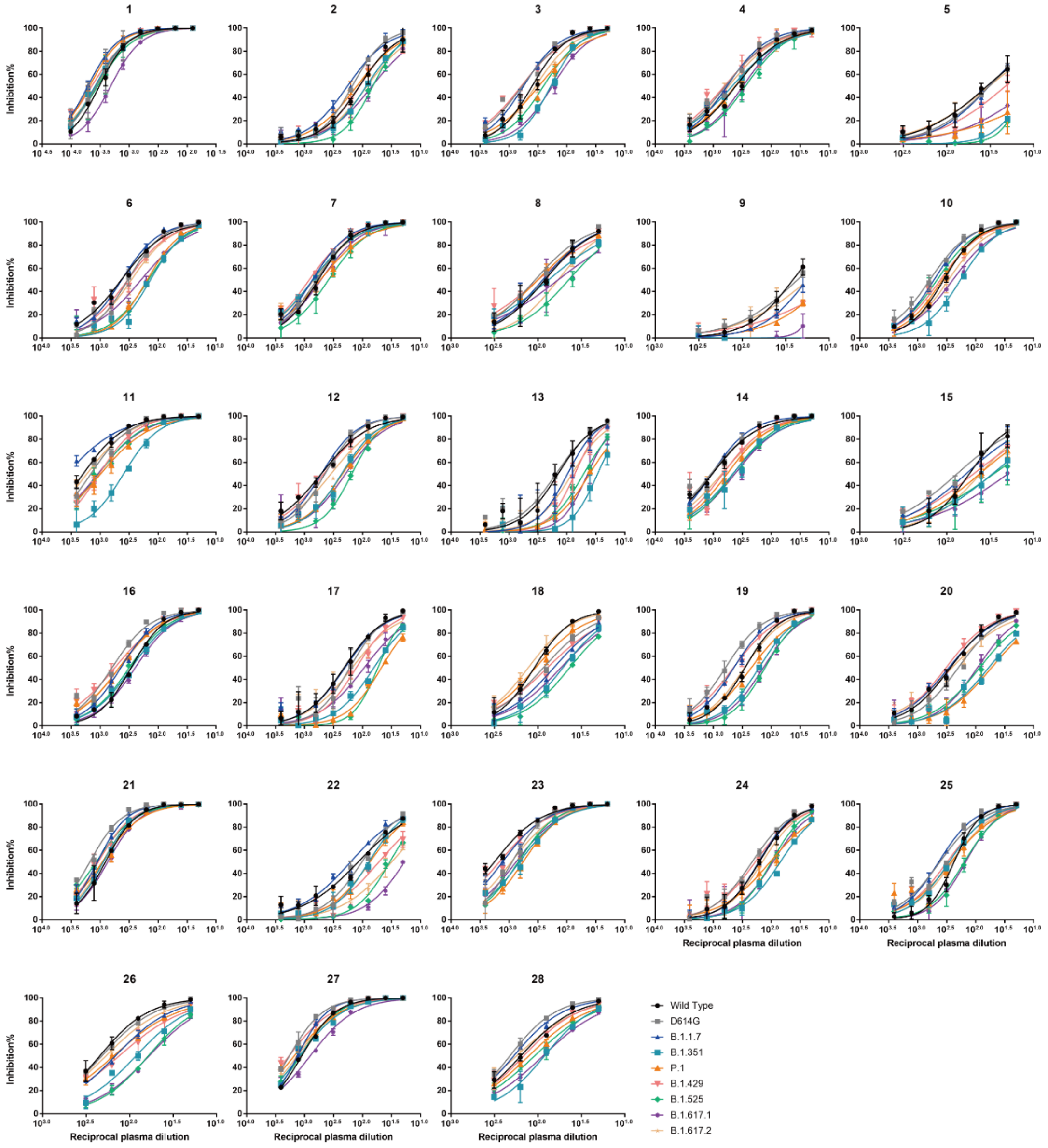

**Figure S2: Neutralization assays of ZF2001 vaccinee plasma against pseudovirus based SARS-CoV-2 variants.** The neutralization assay of plasma from 28 vaccinated volunteers were detected against WT, D614G, B.1.1.7, B.1.351, P.1, B.1.429, B.1.525, B.1.617.1 and B.1.617.2 lineages. Each sample was tested with repeats.

**Table S1. Characteristics of vaccine recipients.**

| Characteristics | Vaccine recipients |
|---|---|
| **No. of participants** | 28 |
| **Age (median, range)** | 30.0 (24-43) |
| **Sex** | |
| Male (%) | 13 (46.4) |
| Female (%) | 15 (53.6) |
| **Time interval between the first and second does (Median day, range)** | |
| 0, 1, 2 Months group (16 persons) | 28.0 (24-29) |
| 0, 1, (4-6) Months group (12 persons) | 27.5 (27-35) |
| **Time interval between the second and third does (Median day, range)** | |
| 0, 1, 2 Months group (16 persons) | 35.0 (35-39) |
| 0, 1, (4-6) Months group (12 persons) | 113.0 (82-137) |
| **Time interval between the third dose and blood sampling (Median day, range)** | |
| 0, 1, 2 Months group (16 persons) | 14.0 (14-14) |
| 0, 1, (4-6) Months group (12 persons) | 64.5 (30-104) |
